# Sequestration of Methane by Symbiotic Deep-Sea Annelids: Ancient and Future Implications of Redefining the Seep Influence

**DOI:** 10.1101/2019.12.23.887653

**Authors:** Shana K. Goffredi, Ekin Tilic, Sean W. Mullin, Katherine S. Dawson, Abigail Keller, Raymond W. Lee, Fabai Wu, Lisa A. Levin, Greg W. Rouse, Erik E. Cordes, Victoria J. Orphan

## Abstract

Deep-sea methane seeps are dynamic sources of greenhouse gas production and unique habitats supporting ocean biodiversity and productivity. Here, we demonstrate new animal-bacterial symbioses fueled by methane, between two undescribed species of annelid (a serpulid *Laminatubus* and sabellid *Bispira*) and distinct methane-oxidizing Methylococcales bacteria. Worm tissue δ^13^C of −44‰ to −58‰ suggested methane-fueled nutrition for both species and shipboard experiments revealed active assimilation of ^13^C-labelled CH_4_ into animal biomass, occurring via engulfment of methanotrophic bacteria across the host epidermal surface. These worms represent a new addition to the few animals known to intimately associate with methane-oxidizing bacteria, and further explain their enigmatic mass occurrence at 150-million-year-old fossil seeps. High-resolution seafloor surveys document significant coverage by these symbioses, beyond typical obligate seep fauna. These findings uncover novel consumers of methane in the deep-sea, and by expanding the known spatial extent of methane seeps, may have important implications for deep-sea conservation.

## Introduction

Methane seeps and hydrothermal vents globally host dense communities of unique organisms, mainly supported via symbioses with chemosynthetic microorganisms, which offer direct access to energy from reduced gases such as H_2_S, H_2_, and CH_4_. Most known symbioses involve sulfide-oxidizing bacteria, while far less common are those that associate with methane-oxidizers (1,2). Nevertheless, many methane-rich seeps exist along continental margins, thus providing the distinct possibility of as-yet unknown symbioses that may act as important sinks for this potent greenhouse gas. As an example, along the Pacific continental margin of northern central Costa Rica, a vast series of seeps occurs (nearly one every 4 km) related to landslide scars, seamount-subduction related fractures, mounds, and faults (3,4). In areas such as this, it is increasingly important to understand the trophic interactions between these ubiquitous seafloor ecosystems and the chemosynthetic animals that they support.

One of the historically recognizable animals to inhabit the periphery of seeps and hydrothermal vents are tube-dwelling ‘fan worm’ annelids placed in Sabellida (including Fabriciidae, Sabellidae and Serpulidae; 5). Several species, in particular, achieve such high densities that they are sometimes referred to as ‘thickets’ (200-400 individuals per m^2^; 6-8). Since the discovery of hydrothermal vents, serpulids have been specifically noted as stable and conspicuous indicators of reduced fluids, forming a virtually continuous zone around hydrothermal vent fields (9). The first serpulid species formally described from hydrothermal vents was *Laminatubus alvini*, discovered along the East Pacific Rise (10). Since then, three serpulid genera (*Laminatubus, Protis*, and *Hyalopomatus*) are known to be common at deep-sea chemosynthetic habitats (11). Closely-related sabellids are usually found in shallow benthic marine habitats, but some have been reported from the deep-sea, and one species (*Bispira wireni*) has been described from hydrothermal vents (12-13). Both serpulids and sabellids have generally been regarded as opportunistic inhabitants in these areas of high productivity, presumably suspension feeding on increased bacteria in the surrounding water, using their ciliated anterior appendages (7,14).

Mass occurrences of serpulids in ancient methane-seep deposits, known as ‘fossil serpulid seeps’, have perplexed scientists for decades and the search for indicator species of previously active paleomethane flux from the seafloor continues to be a high priority for geologists and geobiologists alike. For serpulids, which permanently inhabit calcareous tubes, seven taxa (including *Laminatubus*) have been reported from early Cretaceous to Miocene seep communities ∼ 150-200 MYA (15-17). They appear attached to, or in layers immediately adjacent to, known chemosynthetic vesicomyid clams, lucinid clams, or bathymodiolin mussels, prompting the suggestion that serpulids found in fossil seep deposits were actually regular members of the chemosynthetic community, as opposed to arriving only after seepage had ceased (15). The suggestion that modern day serpulids might be endemic to chemosynthetic habitats was proposed early on by other researchers (7,18), calling into question the assumption of obligate suspension feeding (14). By contrast, annelids within Sabellidae inhabit soft mucous and sediment tubes and thus leave only a minor fossil record (19).

Here, by integrating microbial community profiling, ultrastructural analysis via microscopy, live animal stable isotope tracer experiments, and seafloor surveys, we describe two separate, but closely-related, tube-dwelling annelids that demonstrate active methane-based nutrition via methanotrophic bacterial symbionts. These newly discovered methane-reliant animals are commonly found at seeps, and vents, worldwide and extend the boundaries of the ‘seep’ habitat, a classification that is increasingly important for regulatory and stewardship efforts concerning fisheries and oil drilling in the deep-sea (20).

## Results

At a seep site known as Jaco Scar at 1768-1887 m depth off the west coast of Costa Rica (9°7.1’N, 84°50.4’W) (3,21), a single serpulid species (*Laminatubus* n. sp.) and a single species of sabellid (*Bispira* n. sp.) were abundant in zones of active seepage (Fig. 1). Both annelid species represented a large fraction of the animal community and were observed near and often attached to obligate seep fauna, including *Lamellibrachia barhami* and *Bathymodiolus* spp. mussels. Sampling and shipboard experiments of these two species were carried out with the HOV *Alvin* during RV *Atlantis* expeditions AT37-13 (May-June 2017) and AT42-03 (October-November 2018; Table S1), along with high resolution seafloor surveys by the autonomous underwater vehicle (AUV) *Sentry*, owned and operated by Woods Hole Oceanographic Institution.

**Fig. 1.**
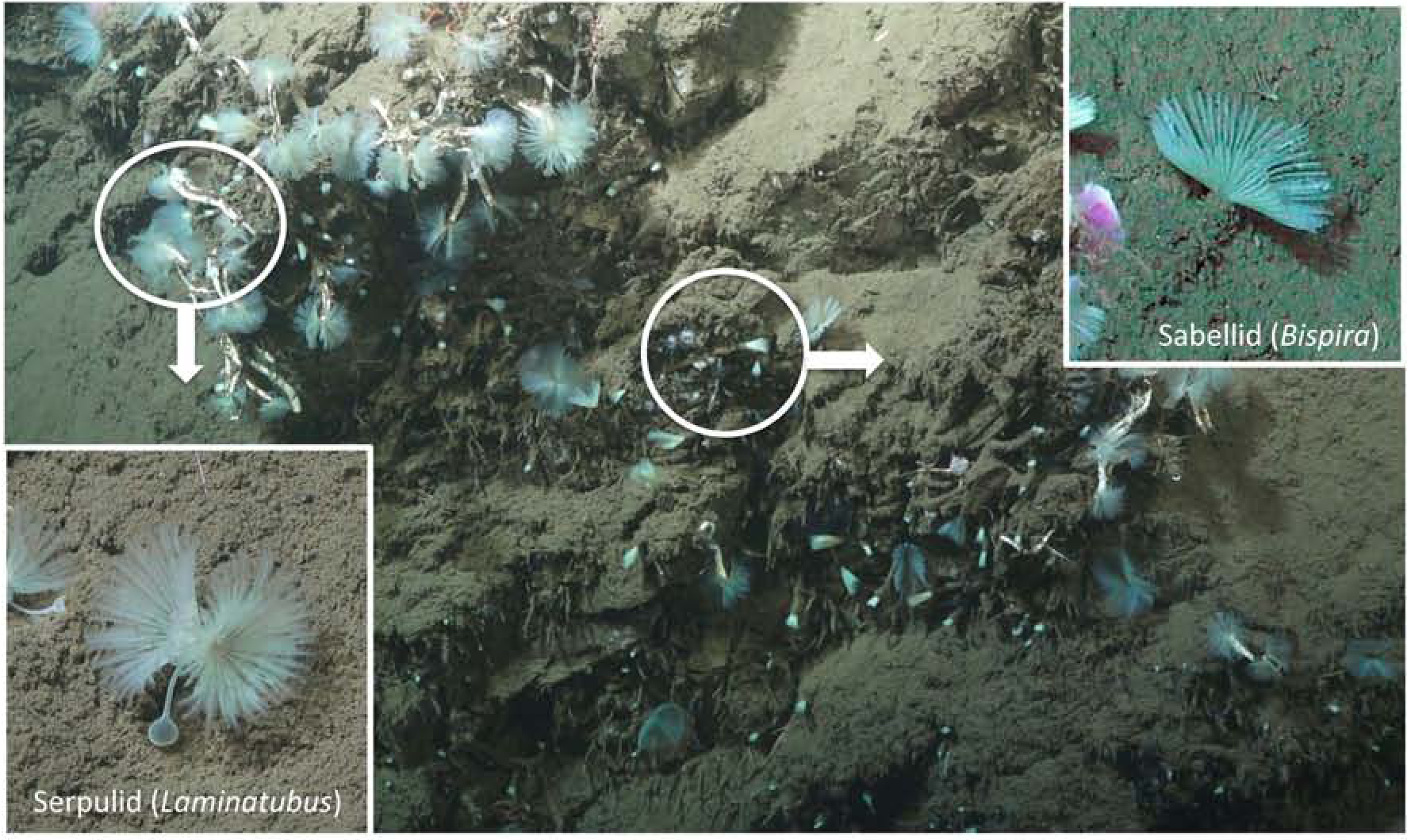
In situ images of the two annelid species featured in this study. A new species of serpulid annelid (*Laminatubus*) and a new species of sabellid (genus *Bispira*), at a site known as Jaco Scar at 1768-1887 m depth (9°7.1’N, 84°50.4’W), along the convergent margin off the west coast of Costa Rica. Insets show the worm species in life position.

### Isotopic evidence of methane-based nutrition

Tissue stable δ^13^C and δ^15^N isotopes of both Jaco Scar *Laminatubus* and *Bispira* species were quite distinct from non-seep members of the same families (Fig. 2). δ^13^C values for the crown appendages, called radioles, of both Jaco Scar worm species were highly negative (−56.9 ± 0.9‰ for *Laminatubus* and −45.3 ± 4.3‰ for *Bispira* (n = 4-6, ± 1 SD)), compared to related known suspension feeders collected from the same general vicinity without active seepage (∼ −18-20‰, n = 5; Fig. 2, Table S1). *Laminatubus* and *Bispira* isotopic values were consistent among all major body parts, including radiolar crown, body wall, and digestive tissues (ANOVA p = 0.65, n = 12 combined for each category; Table S2) and consistent with methane values reported for the water column above Jaco Scar seeps (−50 to −62‰) (21). Particulate organic matter had a δ^13^C of −25‰ in the Jaco Scar seep bottom water (n = 3; Fig. 2).

**Fig. 2.**
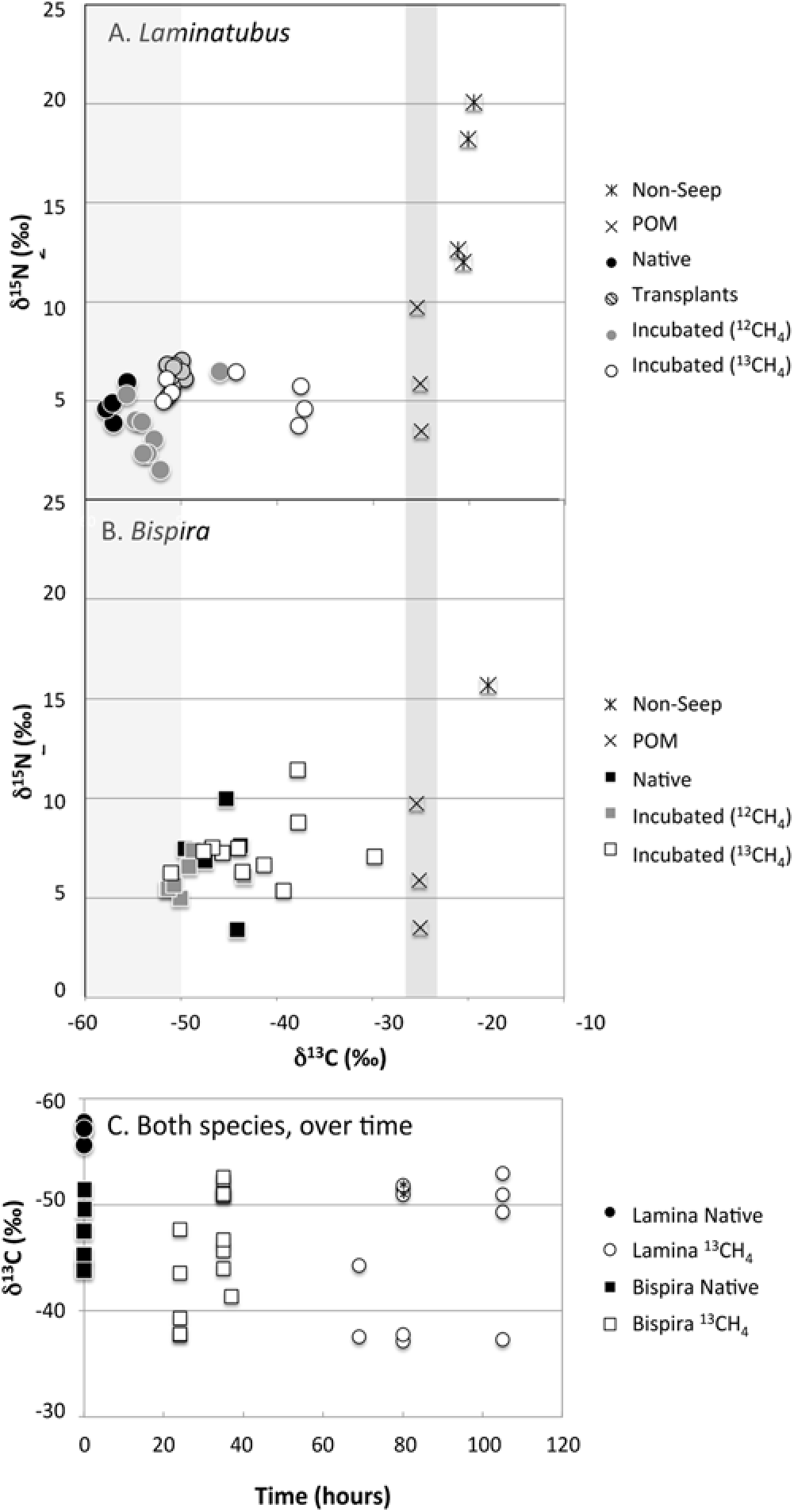
Carbon and nitrogen isotope data for native and experimental animals. Biplots of δ^13^C and δ^15^N values for the crown radioles of (A) the serpulid *Laminatubus* n. sp. and (B) sabellid, *Bispira* n. sp. from the Jaco Scar seep, sampled directly upon collection from the active seep (‘Native’), after an *in situ* transplant for 16 months to an area of no methane venting (“Transplants”), or after shipboard incubation with either 12CH_4_ (“controls”) or ^13^CH_4_ (ex. “^13^CH_4_-incubated”). “Non-seep” sabellids were also collected from an offshore seamount, but were not identified beyond family. The right hand vertical shaded bar highlights the measured values of particulate organic material “POM” (collected from CTD casts over the area of active seepage). The left hand vertical shaded bar indicates the δ^13^C for methane from the water column above Jaco Scar seeps (−62 to −50‰) reported by Mau et al. 2014. (C) Changes in δ^13^C over time are shown for both species. Two ^13^CH_4_-incubated *Laminatubus* that were observed to have a decreased relative abundance of MOX bacteria, based on 16S rRNA analysis (0.3% and 14%; see also Fig. 5) are shown with an asterisk in the symbol.

### Molecular and microscopic evidence of methanotrophic symbioses

In *Laminatubus*, the radioles were arranged in typical serpulid semi-circles, fusing to form a pair of lobes located on each side of the mouth (Fig. 3A). The radioles, which were coiled upon collection, were covered with rows of paired, densely ciliated, pinnules. The dense ciliation, and the bacteria covering the radioles, gave the crown a fuzzy appearance (Fig. 3B). In *Bispira*, the radiolar crown was very long (in some cases as long as the body; crown to body ratio 0.6-1.3; Fig. 4A), with elongate and delicate pinnules arranged along the radioles (Fig. 4B). Of the three tissues associated with *Laminatubus* and *Bispira* in this study, bacteria could only be reliably amplified by PCR from the distal regions of the crown radioles. Bacterial community analysis via 250-bp 16S rRNA barcoding revealed limited diversity, with four putative methanotrophic bacterial OTUs within the Methylococcales Marine Methylotrophic Group 2 (MMG-2; clustered at 99% similarity) dominating the community associated with each annelid species (Fig. 5). MMG-2 OTUs comprised 52-63% of the bacterial community in *Laminatubus* and 72- 90% in native *Bispira* individuals from Jaco Scar, processed immediately following *in situ* collection (= ‘native’). Specific worm-associated MMG-2 OTUs were not detected in either the surrounding water column or nearby sediment (at 0-3 cm depth; data not shown), or in association with two non-seep sabellid specimens collected from nearby inactive areas (Fig. 5; Table S1). This bacterial diversity was remarkably consistent among individuals of both *Laminatubus* and *Bispira* species collected 16 months apart, and from both Jaco Scar and another nearby methane seep known as Mound 12 (Fig. 5; Figure S1; 22). The co-occurring bathymodiolin mussels from Jaco Scar, which are capable of forming methane-based symbioses (23) did not possess MOX bacteria (and had a gill δ^13^C value of −36.6‰, n = 3). Similarly, nearby frenulate siboglinids, also known for methanotrophic symbioses (24), hosted a different MMG-2 OTU that comprised only 15% of their bacterial community (with a δ^13^C value of −39.1‰, n = 3).

**Fig. 3.**
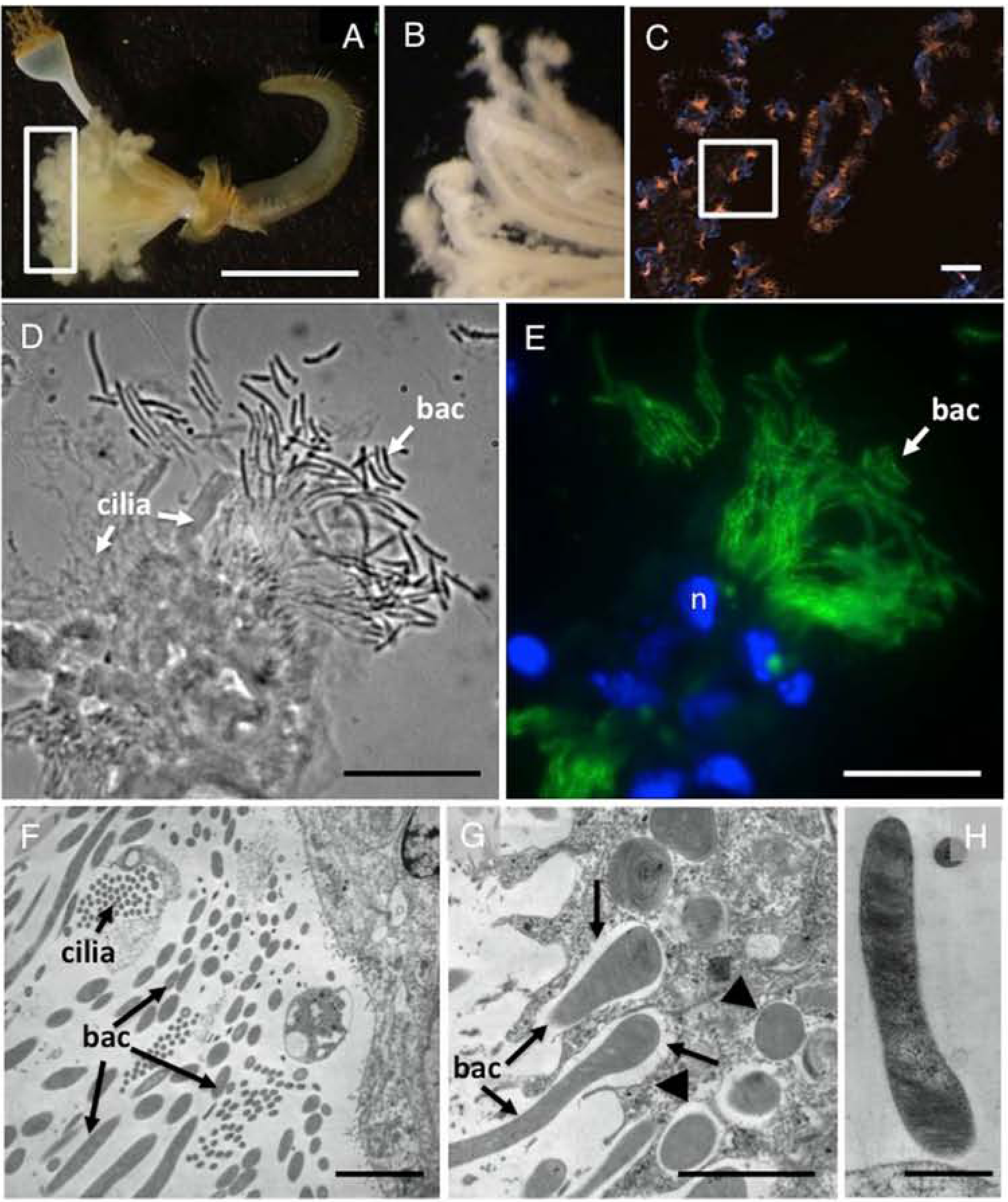
Microscopy of *Laminatubus* n. sp. crown radioles. Light microscopy (A-B), fluorescence (C, E) plus corresponding bright field (D) and transmission electron (F-H) microscopy of *Laminatubus* n. sp. radioles. A FISH probe (MTC851) was designed to be an exact match to the MOX symbionts (related to Methylococcales / Marine Methylotrophic Group 2 - shown orange (C) or green (E)). DAPI-stained nuclei of host cells are shown in blue. TEM images show the MOX bacteria, with dense internal membranes, on the epidermal surface of the host, and also possibly in the process of being engulfed (arrows), and completely engulfed (arrowheads in F and G) by host cells. (H) A close-up of a single MOX filament, attached to the worm epidermis. n, nucleus. bac, bacteria. Scale bars are 1 cm (A), 10 µm (C), 15 µm (D-E), 1 µm (F-H).

**Fig. 4.**
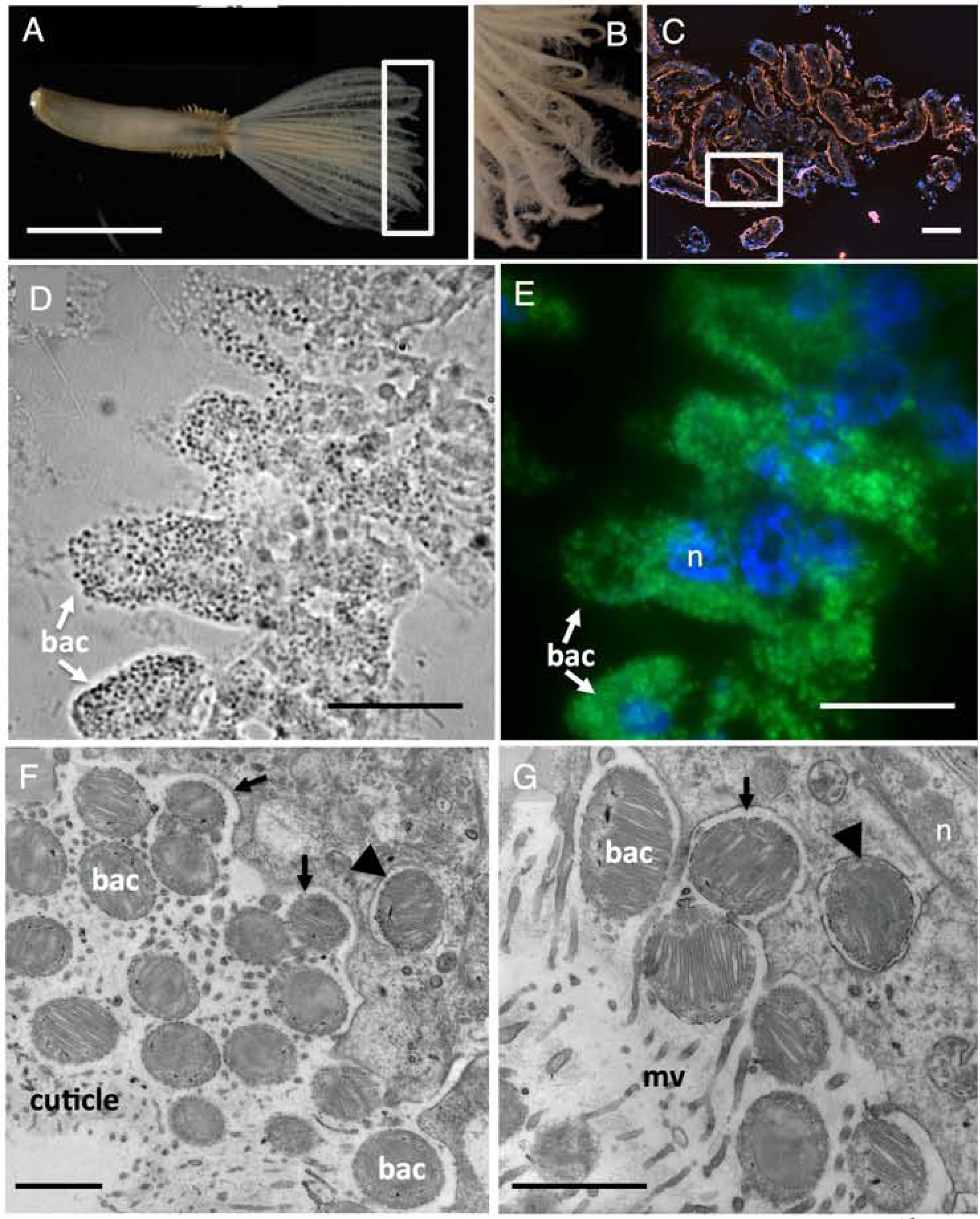
Microscopy of the *Bispira* n. sp. crown radioles. Light microscopy (A-B), fluorescence (C, E) plus corresponding bright field (D) and transmission electron (F-G) microscopy of *Bispira* n sp. radioles. A FISH probe (MTC851) was designed to be an exact match to the MOX symbionts (related to Methylococcales / Marine Methylotrophic Group 2 - shown orange (C) or green (E)). DAPI-stained nuclei of host cells are shown in blue. TEM images show the MOX bacteria, with dense internal membranes, embedded in the cuticle of the host, and also possibly in the process of being engulfed (arrows), and completely engulfed (arrowheads in F and G) by host cells. n, nucleus. mv, microvilli. bac, bacteria. Scale bars are 0.5 cm (A), 10 µm (C), 15 µm (D-E), 1 µm (F-G).

**Fig. 5.**
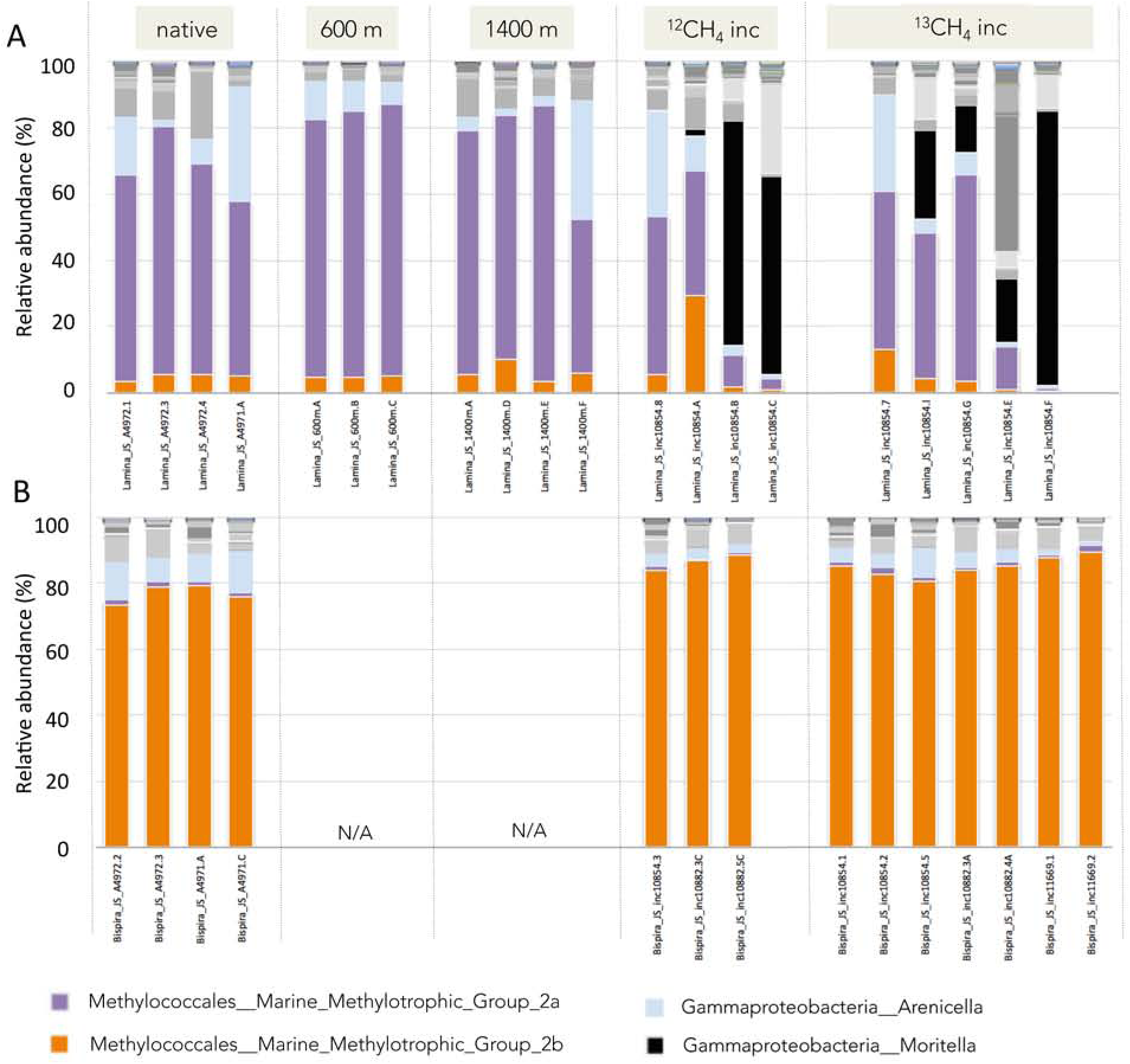
Relative abundance of bacterial phylotypes, based on 16S rRNA. Bacterial community structure for crown radioles of (A) the serpulid *Laminatubus* n. sp. and (B) sabellid *Bispira* n. sp. from Jaco Scar seep, Costa Rica. Tissues were sampled directly upon worm collection from the active seep (‘native’), after transplant for 16 months to areas of very little / no methane venting (‘600 m’ and ‘1400 m’ away) or after shipboard incubation in either ^12^CH_4_ or ^13^CH_4_ (ex. “^13^CH_4_-incubated”). *Bispira* were not transplanted and thus do not have specimens for those categories (‘N/A’). Each color on the graph represents a distinct genus-level phylotype or lowest level available. Phylotypes were grouped to 97% 16S rRNA sequence similarity. Dominant phylotypes are indicated in the key. Genera that were not putative aerobic methanotrophs are shown in gray or black. See DataFile S1 for raw and processed data, as well as representative sequences for all dominant hits.

Targeted PCR amplification and sequencing of the particulate methane monooxygenase gene (pmoA, which performs the oxidation of methane to methanol) and clade-specific 16S rRNA gene was used to better characterize the bacterial methanotrophs in association with *Laminatubus* and *Bispira*. Analysis of a longer region of the 16S rRNA gene, via direct sequencing using a MMG-2 specific reverse primer (MTC850R; based on 25), revealed a single putative methanotrophic 16S rRNA phylotype for each worm species (Fig. S2A). These 16S rRNA phylotypes were 94% similar to each other, and 97% similar to a closest relative (free-living bacteria from Gulf of Mexico seeps and EPR hydrothermal vents, for the *Laminatubus* and *Bispira* phylotypes, respectively. Likewise, a single *pmoA* gene sequence was recovered from each host species, via direct sequencing, with 83% nucleotide similarity between them (Fig. S2B). These same *pmoA* sequences were not detected in the surrounding water column, suggesting that the worms hosted highly specific bacteria, as opposed to those more common in the environment. The worm-associated bacterial *pmoA* sequences, however, did share 99% amino acid identity to *pmoA* genes recovered from other seep environments, including Jaco Scar (26) and notably the frenulate *Siboglinum* from the Gulf of Cadiz (NE Atlantic; 27).

In support of the 16S rRNA-based community structure, abundant bacteria were observed via fluorescence in situ hybridization (FISH) attached to and surrounding the radioles of both *Laminatubus* and *Bispira* from Jaco Scar (Figs. 3 and 4). Via TEM, nearly the entire volume of each putative symbiont cell was devoted to intracytoplasmic membranes (Figs. 3G-H and 4G-I), which are characteristic of MOX bacteria, and the typical site of methane oxidation (28). A Methylococcales - Marine Methylotrophic Group 2 specific FISH probe MTC851 was designed to be an exact match to both MOX symbionts (based on the previously published MTC850 probe). For *Laminatubus*, numerous long (∼10+ μm long × 0.5 μm wide), segmented, filamentous bacteria were observed in rosettes or tufts emerging from the tips of each radiole (Fig. 3C-E). Despite the presence of a second *Arenicella* gammaproteobacteria phylotype (8-35% of the recovered bacterial community, based on 16S rRNA barcoding; Fig. 5), no other bacteria were observed via microscopy on the *Laminatubus* radioles (Figure S3).

Via transmission electron microscopy, numerous filaments with densely packed intracellular membranes were observed directly attached to the epidermis of *Laminatubus* (Fig. 3F-H). For *Bispira*, numerous small (∼0.6 μm), membranous, cocci-shaped bacterial cells (Fig. 4C-E) were embedded in an extracellular collagenous cuticle, penetrated by microvilli (Fig. 4F). TEM analysis of both species showed that MOX bacteria were not only attached to the epidermal surface of the worms but were clearly in the process of engulfment by host tissue (Figs. 3G, 4G). Bacterial cells with compacted and disorganized membranes appeared deep in the worm tissues, surrounded by intracellular structures interpreted as digestive vacuoles (Figure S4D-G).

Further, hybridization with the lipophilic dye FM4-64 revealed lipid-rich, presumably host-sourced, structures near the sites of bacterial attachment (Figure S4B-C). Examination of *Bispira* revealed a cuticle matrix comprised primarily of mannose, as stained by *Hippeastrum* Hybrid Amaryllis lectin (HHA; Figure S4A). Mannose is a constituent of the cuticle in some annelids (29) and is the lock-and-key sugar used in other symbioses (30). In agreement with molecular analysis, fluorescence microscopy did not reveal obvious bacteria in the digestive system for either species (Figure S5).

### In Situ Transplantation of methanotroph-bearing serpulids

*Laminatubus* specimens were transplanted by submersible *in situ* from an area of active seepage to two inactive muddy areas 600-1400 m away from the active site, in the general direction of methane plume flow (south-southwest, based on deployed current meters). These worms survived for 16 months and those examined had retained their respective MOX bacterial phylotypes (which comprised 73-83% of the bacterial community, based on 16S rRNA amplicon sequencing; Fig. 5) and light tissue δ^13^C values (−50.6 ± 0.7‰; Fig. 2A). Tissue δ^13^C values of the transplanted worms, however, were slightly heavier than the δ^13^C values of individuals that remained in the active seep area (ANOVA P < 0.00001; Fig. 2A; Table S2).

### ^13^CH_4_ tracer evidence of active methane incorporation by host worms

In order to examine the possibility of methane oxidation and assimilation of methane-derived carbon into host tissues, whole worms with intact symbionts were incubated for 24-105 hours in the presence of ^13^C-labelled CH_4_ (Table S3). The generation of substantial ^13^C-labelled dissolved inorganic carbon in the surrounding seawater was measured within 15 hours of incubation, confirming CH_4_ conversion to CO_2_ (Figure S6). This included a set of incubations where live worms were removed from their tubes prior to incubation in order to minimize the potential for methanotrophic activity associated with bacteria on the tube surface. Within 24 hours of incubation, clear assimilation of 13C-labelled CH_4_ was observed within animal tissues (Fig. 2). For *Laminatubus*, δ^13^C values increased from −52.9 ± 3.0‰ (n = 8; for control worms exposed to ^12^CH_4_) to −44.4 ± 6.9‰ for ^13^CH_4_-incubated animals (n = 7; ANOVA P = 0.012, F = 7.847), with the tissues of three individuals revealing a shift to ∼ −37‰ δ^13^C (Fig. 2). Similarly, ^13^CH_4_-incubated *Bispira* had heavier δ^13^C values (−42.2 ± 5.9‰, n = 11), compared to control animals (−48.9 ± 2.9‰, n = 6; ANOVA P = 0.021, F = 6.6471), again with four individuals showing δ^13^C values as heavy as −29 to −39‰ within 37 hours of shipboard ^13^CH_4_ exposure (Fig. 2).

The variable response of ^13^CH_4_ incorporation by some *Laminatubus* individuals (Fig. 2), however, could be explained by the variable community representation of methane-oxidizing bacteria observed for ^13^CH_4_-incubated *Laminatubus* (0.3-44%; Fig. 5). For example, several *Laminatubus* that did not take up as much ^13^CH_4_ (δ^13^C values ∼ −52‰; Fig. 2A) were found to be overgrown during the incubation period by *Moritella* (ex. up to 61-84%, based on 16S rRNA sequencing; Fig. 5), a bacterium not observed in native worms. FISH microscopy confirmed the lack of methane-oxidizing bacteria on the crowns of these individuals. All of the *Laminatubus* and *Bispira* collected in 2017 and 2018 hosted abundant MOX bacteria, thus we do not think this reflects a symbiosis that is sensitive to environmental conditions, but rather an artifact caused by containment during experimental incubations, especially after removal of some from their calcareous tubes. Notably, individuals that showed the greatest ^13^CH_4_ assimilation (a positive δ^13^C shift of ∼ 16‰) maintained a dominance of the MOX bacterial phylotype throughout the incubations (∼44% MOX bacteria; Fig. 5).

### Mapping serpulid community scale at the Jaco Scar seep

Using the autonomous underwater vehicle *Sentry*, the presence of serpulids, with their easily observed white calcareous tubes, was mapped at the Jaco Scar seep region (Fig. 6). AUV *Sentry* travelled a total of ∼44 km in a grid pattern (during 4 dives in 2017 and 2018), at an altitude of ∼7m, taking high-resolution digital georeferenced photographs (∼1 image every 3 seconds). In photos with clear visibility, the presence or absence of carbonate hard substrate and obligate seep organisms — bacterial mats, vestimentiferan tubeworms, vesicomyid clams, bathymodiolin mussels — was noted. These species and substrate occurrences were resampled into 5 m grid cells in the survey area and mapped over the corresponding multibeam bathymetric data, using ArcGIS. Serpulids were categorized as “seep-associated” if obligate seep organisms were observed in the same grid cell. This autonomous approach revealed the seascape-scale distribution of these organisms, as compared to the limited range of observations that can be made visually from an ROV or human-occupied vehicle. Overall, serpulids were observed to cover at least 65,000 m^2^ at the Jaco Scar seep site and importantly extended the boundary, in some areas, ∼300 m beyond the observed obligate seep organisms (Fig. 6). In particular, the distribution of serpulids extended the furthest to the southwest, which was the prevailing direction of water current flow from the active seep during the study period from 2017-2018. Only a single species of serpulid was observed at the Jaco Scar seep site, thus it was assumed that the serpulids visualized by *Sentry* were all *Laminatubus*. Similar analyses were not possible for *Bispira*, since they are not obvious in *Sentry* images, but they were most often found co-occurring with *Laminatubus*.

**Fig. 6.**
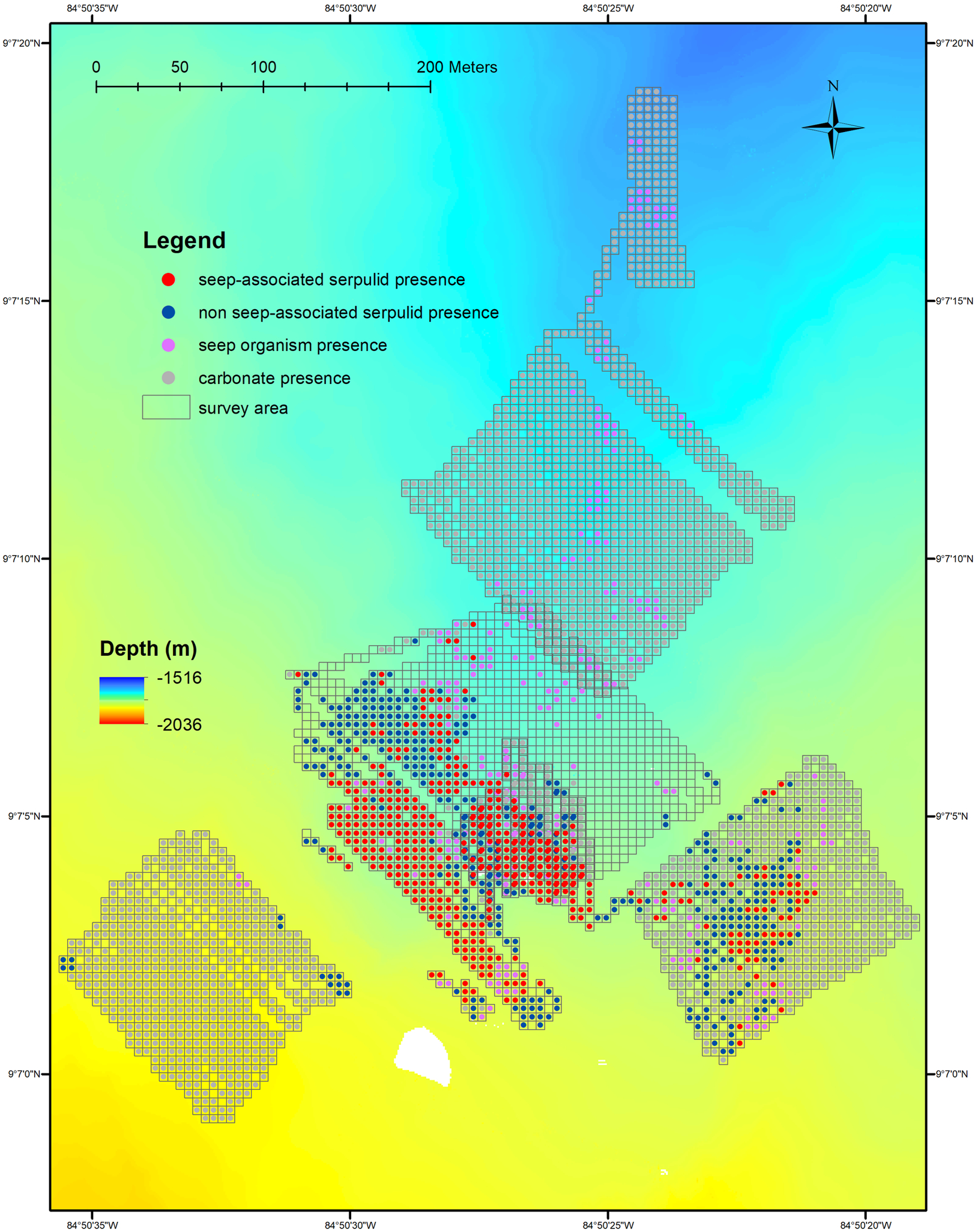
Bathymetric map of the Jaco Scar seep region, as surveyed by the AUV *Sentry*. Location of the Jaco Scar seep study site, off the west coast of Costa Rica, and the 44 km track surveyed by AUV *Sentry*, equipped with high-resolution mapping tools (sidescan sonar, multibeam echo-sounder, digital still camera), traversed during four dives in 2017 and 2018. Moving at an average speed of 0.54 m/s, *Sentry* traveled at an average altitude of 6.9 m, and took ∼ 20 photos / min. Out of a total of 42,272 downward-facing photos, 28,962 photos were annotated with serpulids. Serpulid presence was categorized by the occurrence of other obligate seep fauna within the same 5 meter grid cell (“seep-associated”) and the absence of other obligate seep fauna within the same grid cell (“non seep-associated”). The distribution of seep fauna and carbonate rock without serpulids is also shown.

## Discussion

Deep-sea cold seeps are dynamic point sources for methane in the environment. Most studies in these habitats have emphasized environmental methane-oxidizing bacteria and archaea as important methane sinks; however, certain animals can also enrich for and host dense communities of methanotrophs. Here, we demonstrate active methane consumption by two new species of annelid (a serpulid *Laminatubus* sp. and sabellid *Bispira* sp.) and distinct methane-oxidizing gamma-proteobacteria within the Marine Methylotrophic Group 2 (Methylococcales).

In both modern and paleo-ecosystems, isotope analysis is frequently used as an indicator of reduced chemical utilization, whether via methane or sulfide oxidation coupled to carbon fixation. The highly negative δ^13^C values in *Laminatubus* and *Bispira* tissues (−44‰ to −58‰) suggest a significant contribution of methane-derived carbon to their biomass. Two possibilities for the relatively light animal δ^13^C values involve aerobic methane-oxidizing (MOX) bacteria, which occur ubiquitously in areas with exposure to both oxygen and methane, and have been described from the water column near Jaco Scar and other seep areas (26, 28). Either the worms selectively filter feed on MOX bacteria from the overlying water or they form specific methane-based symbioses with MOX bacteria. Prior evidence exists for selective feeding on aerobic methanotrophs by annelids in seeps, particularly sediment dwelling ampharetids that form crater-like depressions in the sediments as they errantly deposit feed (31-32). For *Laminatubus* and *Bispira*, however, selective feeding does not seem likely given that water column methanotrophs are only a small fraction of the total bacteria (4,33), and therefore presumably unable to support the very high biomass of these worms, which cannot leave their tubes. Furthermore, the δ^13^C of Jaco Scar POM from the surrounding water (−25‰) was not reflected in the tissues of either species, and we found no evidence for bacteria in their digestive systems. Alternately, aerobic MOX bacteria are known to live in intimate symbioses with marine invertebrates, including siboglinid tubeworms, bathymodiolin mussels, provannid snails, and cladorhizid sponges (reviewed in 2). *Bathymodiolus* mussels possess MOX bacteria intracellularly in gill tissues, with corresponding negative δ^13^C values (−40 to −80‰) (2,34) — a similar resemblance of *Laminatubus* and *Bispira* tissues to water column CH_4_ values from Jaco Scar (−50 to −62‰; 21).

Molecular evidence supported a specific and persistent symbiotic association between both *Laminatubus* and *Bispira* and distinct MOX bacteria within the Methylococcales MMG-2 group. These particular bacteria were present in individuals collected 16 months apart, and conversely not detected in the surrounding water column, underlying sediments, or in association with closely-related worms from nearby inactive areas. Microscopy was striking in that abundant long filamentous MOX bacteria covered the radioles of *Laminatubus* while numerous smaller cocci-shaped bacteria were embedded in the collagenous cuticle of *Bispira* radioles. The Marine Methylotrophic Group 2 has recently been shown to associate with invertebrates, including asphalt seep sponges (35) and *Siboglinum* worms (27). Interestingly, the bathymodiolin mussels, which are known to form symbioses with MOX bacteria, have not done so at the Jaco Scar methane seep sites. Thus, it appears that serpulids and sabellids have assumed an important symbiotic niche enabling the exploitation of methane for animal nutrition.

Transplanted *Laminatubus* survived for 16 months and those examined retained their respective MOX bacterial phylotypes and had light tissue δ^13^C values suggestive of persistent CH_4_ metabolism and assimilation. *Bathymodiolus* mussels with methanotrophic symbionts have likewise been shown to assimilate methane concentrations far below detection limits (36), but in the case of *Laminatubus*, the affinity for methane by the methanotrophic symbionts is not yet known. Tissue δ^13^C values of the transplanted worms, however, were significantly heavier than individuals that remained in the active seep area, perhaps indicating breakdown of tissue biomass (the transplanted worms appeared more translucent in nature), a lack of carbon fractionation during starvation, or increased suspension feeding activity, which has also been observed in mixotrophic *Bathymodiolus* mussels (37).

Whole worms with intact symbionts, incubated in the presence of ^13^C-labelled CH_4,_ revealed active oxidation of CH_4_ and assimilation into host tissues. This appears to be the first example of marine invertebrates hosting a nutritional methane-oxidizing bacterial symbiont externally. This raises the important question of how the worms acquire carbon, or other nutrients, from their bacterial symbionts. Typically, organic carbon is passed from symbiont to host by either translocation of small organic molecules or by host intracellular digestion of the symbionts (38). The distinction between these two mechanisms in methanotrophic symbioses is often determined by ultrastructural observations and incubations with isotopically-labelled methane. In bathymodiolin mussels, a time frame of carbon transfer to symbiont-free tissues on the order of hours has been attributed to translocation, whereas transfer in 1-5 days, consistent with the time frame of our experiments, is indicative of digestion (38). TEM analysis of both annelid species showed that MOX bacteria were not only attached to the epidermal surface of the worms but were clearly in the process of engulfment by host tissue. Bacterial cells with compacted and disorganized membranes appeared deep in the worm tissues, surrounded by intracellular structures interpreted as digestive vacuoles. Further, hybridization with a lipophilic dye revealed lipid-rich structures near the sites of bacterial attachment, supporting internalization and possible host endocytosis of bacteria (39). To our knowledge, the only other example of this mechanism for nutrient acquisition is in thyasirid clams, whose extracellular symbionts are periodically engulfed and digested by host gill epithelial cells (40). It remains a possibility that *Laminatubus* and *Bispira* also recover specific small carbon molecules from the bacteria (e.g. fructose-6-phosphate, resulting from the ribulose monophosphate pathway, likely used by this group of gammaproteobacterial MOX bacteria for assimilation of formaldehyde, following the oxidation of methane). Follow-up investigations with live specimens are required to determine the specific mechanism for carbon exchange and exactly how much of the nutrition of both annelids is supported by methane.

## Conclusion

Two species of tube-dwelling annelids from within Sabellida have independently established highly-specific, persistent, nutritional symbioses with two closely-related aerobic methanotrophic bacteria. The assumption, until now, was that most sabellids and serpulids acquire nutrients via suspension feeding (14). We are reminded that heterotrophy involving suspension feeding was also initially proposed for the giant hydrothermal vent tubeworm *Riftia pachyptila*, which we now know relies entirely on sulfide-oxidizing chemoautotrophic symbionts for nutrition (41-43). Very little is known about the ecology of extant serpulids and sabellids in chemosynthetic ecosystems, and many species reported from hydrothermal vents and cold seeps remain identified only to the family level (ex. 18). *Laminatubus*, in particular, is known from other seeps (e.g. Pescadero Transform fault) and vents (e.g. Alarcon Rise; 8), and additional serpulids have been observed at hydrothermal vents with significant methane values in end member fluids, including Logatchev, Rainbow, Menez Gwen and Lucky Strike (44). Thus, the occurrence of methanotrophic symbionts associated with these ubiquitous animals is likely to extend beyond the Costa Rica margin seeps and may even extend beyond species of *Laminatubus* and *Bispira*. Further, these results also explain the unusually high abundance of serpulids at ancient hydrocarbon seeps worldwide (up to ∼50% of estimated total animal biomass; 17)

Using the autonomous underwater vehicle *Sentry*, the presence of serpulids was mapped at the Jaco Scar seep region. Despite the power of this approach, to our knowledge, only one other publication has similarly employed an autonomous vehicle for high resolution spatial characterization of cold seep communities (45). Overall, serpulids were observed to cover at least 65,000 m^2^ at the Jaco Scar seep site and importantly extended the boundary, in some areas, ∼300 m beyond the observed obligate seep organisms (e.g. clams and siboglinid tubeworms). The general expanse of these worms beyond obvious observable vent or seep features is in agreement with past studies (6-7) and provides ecological support for the concept that they significantly widen the influence of methane in deep-sea ecosystems. Given their expansive presence at seeps along the Costa Rica margin, and high densities at other known vent and seep sites, *Laminatubus* and *Bispira* represent a potentially valuable and previously unrecognized ecosystem service – that is the oxidation and sequestration of methane, a potent greenhouse gas. We have yet to quantify the extent of the ecosystem services provided by these symbioses, but our observations appear to expand the ‘sphere of influence’ (sensu 46) of methane in the deep-sea, including the transfer of methane-based carbon across areas of high seepage to the periphery of seeps and into the background deep sea.

What might it mean for the ocean to lose these ecosystem services? Continental margins at bathyal depths, where methane seeps typically occur, are increasingly being exploited (47). Seeps, as essential fish habitats, contribute to the productivity of bottom fisheries (48) and seeps in general often co-occur with areas of interest to the oil and gas industry (49). Current national and international spatial management prohibits particular activities from certain areas, for example exclusion zones where sensitive species or habitats, including cold seeps, are present (49). This, of course, depends on an informed understanding of how to locate and define the seep ecosystem footprint, not just via the presence of methane bubbles and traditionally large visible, symbiont-bearing animals. The boundaries between seep ecosystems and the surrounding deep sea are not as sharp in space or time as often depicted (46). A new recognition of abundant methanotrophic serpulid and sabellid assemblages as an expansion of the traditional notion of the methane seep ecosystem brings us closer to accurate and knowledgeable use of the spatial influence of seepage to guide regulatory actions along the world’s active continental margins.

## Materials and Methods

### Sampling

All *Laminatubus* and *Bispira* specimens were collected from active seep sites with the HOV *Alvin* during RV *Atlantis* expeditions AT37-13 (May-June 2017) and AT42-03 (October-November 2018) from Jaco Scar (∼ 1800 m depth) and Mound 12 (∼ 1000m depth), off the coast of Costa Rica (Table S1). Targeted water samples (2-5 L) were collected via Niskin bottle, also via HOV *Alvin. Laminatubus* specimens were transplanted, via HOV *Alvin* (during RV *Atlantis* expedition AT37-13) to two separate areas 600 m and 1400 m from the sites of active venting at Jaco Scar. They were recovered, via HOV *Alvin* (during RV *Atlantis* expedition AT42-03, 16 months later. Additional non-seep species were collected during ROV *SubBastian* dives SO224 and SO227 (in January 2019) near sites known as Coco South and Seamount 6 (320-530 m depth; Table S1). Upon recovery immediately following each dive, worms were subsampled for various analyses or shipboard experiments.

### Isotope Analysis

Tissue samples were dissected at sea, rinsed in milli-Q water, and frozen at −20°C until shipment to the stable isotope laboratory (at Washington State University). Carbon and nitrogen isotope values of worm tissues and particulate organic carbon, 4L of water pulled onto glass fiber filters, were determined by isotope ratio mass spectrometry. Freeze dried samples (0.3-0.7 mg dry weight) were packaged in tin capsules for mass spectrometry, and analyzed using a Costech (Valencia, CA USA) elemental analyzer interfaced with a continuous flow GV instruments (Manchester, UK) Isoprime isotope ratio mass spectrometer (EA-IRMS) for ^15^N/^14^N and ^13^C/^12^C ratios. Measurements are reported in δ notation [per mil (‰) units] and a protein hydrolysate calibrated to NIST reference materials was used as a standard. Precision for δ^13^C and δ^15^N was generally ± 0.2‰ and ± 0.4 ‰, respectively.

### Molecular Analysis

Specimens for molecular analysis were preserved either immediately upon collection, or immediately following shipboard experiments, in ∼90% ethanol and stored at 4°C. Total genomic DNA was extracted from tissues (50-130 mg) using the Qiagen DNeasy kit (Qiagen, Valencia, CA, USA) according to the manufacturer’s instructions. Water samples taken by HOV *Alvin* were filtered (2L) onto a 0.22 µm Sterivex-GP polyethersulfone filter (Millipore-Sigma, St. Louis, MO, USA) and frozen at −80°C until DNA analysis. DNA extraction was also performed using the Qiagen DNeasy kit, according to the manufacturer’s instructions, with the exception of the first step where 2 ml of ATL lysis buffer was added to the sterivex filter, via luer lock and syringe, and rotated at 56°C for 12 hours. This solution was recovered from the filter, also via luer lock and syringe, and processed as usual. A 478-bp region of the gene coding for *pmoA* (subunit of particulate methane monooxygenase enzyme) was amplified using the primers pmoA189f (5′-GGNGACTGGGACTTCTGG-3′) and pmoA661r (5′-CCGGMGCA-ACGTCYTTACC-3) (50). A 823-bp fragment of the 16S rRNA gene was amplified using the primers 27F and MTC850R, originally designed as a FISH probe targeting the MMG-2 group (41). Annealing conditions of 56°C and 54°C were used for *pmoA* and 16SrRNA, respectively, for 25 cycles. Amplification products were sequenced directly using Sanger sequencing, via Laragen Inc., and submitted to GenBank (accession numbers TBD - submitted July 24, 2019). Close environmental and cultured relatives were chosen using top hits based on BLAST (www.ncbi.nlm.nih.gov).

The V4-V5 region of the 16S rRNA gene was amplified using archaeal/bacterial primers with Illumina (San Diego, CA, USA) adapters on 5’ end 515F (5′-TCGTCGGC-AGCGTCAGATGTGTATAAGAGACAGGTGCCAGCMGCCGCGGTAA-3′) and 806R (5′-GTCTCGTGGGCTCGGAGATGTGTATAAGAGACAGGGACTACHV-GGGTWTCTAAT-3′) (51). PCR reaction mix was set up in duplicate for each sample with Q5 Hot Start High-Fidelity 2x Master Mix (New England Biolabs, Ipswich, MA, USA) with annealing conditions of 54°C for 25 cycles. Duplicate PCR samples were then pooled and 2.5 µL of each product was barcoded with Illumina NexteraXT index 2 Primers that include unique 8-bp barcodes (P5 5’-AATGATACGGCGACCACCGAG-ATCTACAC-XXXXXXXX-TCGTCGGCAGCGTC-3’ and P7 5’-CAAGCAGAA-GACGGCATACGAGAT-XXXXXXXX-GTCTCGTGGGCTCGG-3’). Amplification with barcoded primers used conditions of 66°C annealing temperature and 10 cycles. Products were purified using Millipore-Sigma (St. Louis, MO, USA) MultiScreen Plate MSNU03010 with vacuum manifold and quantified using Thermo-Fisher Scientific (Waltham, MA, USA) QuantIT PicoGreen dsDNA Assay Kit P11496 on the BioRad CFX96 Touch Real-Time PCR Detection System. Barcoded samples were combined in equimolar amounts into single tube and purified with Qiagen PCR Purification Kit 28104 before submission to Laragen (Culver City, CA, USA) for 2 × 250bp paired end analysis on the Illumina MiSeq platform with PhiX addition of 15-20%.

MiSeq 16S rRNA sequence data was processed in Quantitative Insights Into Microbial Ecology (v1.8.0). Raw sequence pairs were joined and quality-trimmed using the default parameters in QIIME. Sequences were clustered into *de novo* operational taxonomic units (OTUs) with 99% similarity using UCLUST open reference clustering protocol, and then, the most abundant sequence was chosen as representative for each *de novo* OTU. Taxonomic identification for each representative sequence was assigned using the Silva-119 database, clustered at 99% similarity. A threshold filter was used to remove any OTU that occurred below 0.01% in the combined samples dataset. The sequence data was further rarified by random subsampling to equal the sample with the least amount of sequence data (4006 sequences). The raw and processed sequence data, as well as representative sequences, are available in DataFile S1.

### Fluorescence Microscopy

Specimens for fluorescence in situ hybridization (FISH) microscopy were initially preserved in 4% sucrose-buffered paraformaldehyde (PFA) and kept at 4°C. These PFA-preserved specimens were rinsed with 2× PBS, transferred to 70% ethanol, and stored at –20°C. Tissues were dissected and embedded in Steedman’s wax (1 part cetyl alcohol: 9 parts polyethylene glycol (400) distearate, mixed at 60°C). An ethanol: wax gradient of 3:1, 2:1 and 1:1, and eventually full strength wax, was used to embed the samples (1 h each treatment). Embedded samples were sectioned at 2-5 µm thickness using a Leica RM2125 microtome and placed on Superfrost Plus slides. Sections were dewaxed in 100% ethanol rinses. Hybridization buffers and wash buffers were prepared according to 52, using 35% formamide in the hybridization buffer and 450 mM NaCl in the wash solution, and fluorescent probes were at a final concentration of 5 µg/ml. Initially, we used the MTC850 probe (5′-ACGTTAGC-TCCGCCACTA-3′; labelled with Cy3 Figure 3C/4C), designed to target Marine Methylotrophic Group (MMG) 2 MOX, with 1-bp mismatch to the *Laminatubus* and *Bispira* MOX symbionts (31). Ultimately, a probe that was an exact match to both serpulid and sabellid MOX symbionts (MTC851; 5′-ATACGTTAGCTCC-ACCAC-T-3′ labelled with FITC; Figure 3E/4E), was designed, and importantly had 5-bp mismatches to *Arenicella*, the other bacterial phylotype recovered via amplicon sequencing from *Laminatubus*. A universal bacterial probe mix (Eub338 .I-III) (53), was also used. Probes were hybridized at 46°C for 4-8 h, followed by a 15 min wash at 48°C. To visualize the extracellular polymer structure, the sections were stained for 15 minutes with the mannose-specific lectin stain Hippeastrum Hybrid Amaryllis (HHA) labelled with TRITC, at a final concentration of 100 µg/ml in PBS, and then rinsed for 10 minutes with water. To investigate endocytosis, the sections were stained for 15 minutes with a lipophilic FM4-64 dye at a final concentration of 10 ug/ml in PBS and then rinsed for 10 minutes with water. Sections were counterstained with 4’6’-diamidino-2-phenylindole (DAPI, 5 mg/mL) for 1 min, rinsed and mounted in either Citifluor or Vectorshield. Tissues were examined by epifluorescence microscopy using either a Nikon E80i epifluorescence microscope with a Nikon DS-Qi1Mc high-sensitivity monochrome digital camera or a Zeiss Elyra microscope with an ANDOR-iXon EMCCD camera.

### Transmission Electron Microscopy

For transmission electron microscopy, radioles of several individuals of each species were fixed in 2.5 % glutaraldehyde buffered in 0.05 M phosphate buffer with 0.3 M NaCl (2-24 h at 4°C). Ruthenium red (∼0.5%) was added to the fixative. The specimens were rinsed in the same buffer and postfixed with 1% OsO_4_ for 30 min. The radioles were dehydrated in an ascending alcohol series and embedded in Spurr’s resin. Silver-interference colored sections (65–70 nm) were prepared using a diamond knife (Diatome Histo Jumbo) on a Leica Ultracut S ultramicrotome. The sections were placed on Formvar-covered, single-slot copper grids and stained with 2% uranyl acetate and lead citrate in an automated TEM stainer (QG-3100, Boeckeler Instruments). The sections were examined using a Zeiss EM10 transmission electron microscope with digital imaging plates (Ditabis Digital Biomedical Imaging Systems, Germany).

### Shipboard Isotope Labeling Experiments and Analysis

In order to assess interactions between the methanotrophic bacteria and host annelids, short-term stable isotope incubation experiments with ^13^C-labeled methane (^13^CH_4_) were set up at sea using either *Laminatubus* or *Bispira* individuals removed from tubes (to avoid activity caused by bacteria transiently associated with the tubes), or worm-colonized rocks from the Jaco Scar seep. Unlabeled control incubations were included in each incubation series. All treatments were incubated at 4°C in sealed mylar bags with 0.2-µm filtered bottom seawater from the collection site. Methane was added to each incubation to represent approximately one quarter of the total headspace. For one series of incubations, sterile filtered air was added at 15 and 37 hours incubation time. Initial conditions for all incubations, assuming equilibrium between gases and seawater and no starting methane in the filtered seawater, are displayed in Table 3. At the end of each incubation, worms were picked from the rocks, tissues were dissected, rinsed in milli-Q water and frozen −20°C until analysis, according to the Isotopic analyses section above. Additionally, water samples for dissolved inorganic carbon (DIC) measurements were filtered through a 0.2 µm PES filter into helium (He) flushed, 12-ml exetainer vials (Labco Ltd, Lampeter, UK), following the addition of 100 µl ∼40% phosphoric acid. Samples were stored upright at room temperature. Vials were sampled using a GC-PAL autosampler (CTC Analytics, Zwingen, Switzerland) equipped with a double-holed needle that transferred headspace using a 0.5 ml min^-1^ continuous flow of He to a 50 µm sample loop prior to separation by a PoraPlotQ fused silica column (25m; i.d. 0.32 mm) at 72°C. CO_2_ was then introduced to a Delta V Plus IRMS using a ConFlo IV interface (Thermo Scientific, Bremen, Germany) in the Caltech Stable Isotope Facility. A sample run consisted of 3 reference CO_2_ gas peaks, 10 replicate sample injections, and 2 final reference CO_2_ peaks. δ^13^C values were corrected for sample size dependency and then normalized to the VPDB scale with a two-point calibration. using NBS-19 and a previously calibrated laboratory carbonate as internal standards. The overall precision of this technique is estimated to be better than +/- 0.2‰, based on multiple analysis of independent standards as samples.

### Seafloor Surveys with the Autonomous Underwater Vehicle Sentry

In the current study, we used WHOI’s autonomous underwater vehicle (AUV) *Sentry*, equipped with sidescan sonar, a Reson 7125 multibeam echosounder, and a down-looking digital color camera, to collect bathymetric and photographic data. In 2017 and 2018, AUV *Sentry* was deployed off of *R/V Atlantis*, during four dives at Jaco Scar, analyzed in this study. In the photo survey portion of each dive, *AUV Sentry* moved back and forth on tightly spaced tracks (i.e. “mowing the lawn”) following a preset navigation plan. Moving at an average speed of 0.54 m/s, *Sentry* traveled a total of 44.10 km across four dives (#433, 436, 501, 503), at an average altitude of 6.92 m, and took ∼ 20 photos / min. Out of a total of 42,272 downward-facing photos, 28,962 photos were annotated with serpulids. For photos with clear visibility, the presence or absence of seep foundation species—bacterial mats, *vestimentiferan* tubeworms, *vesicomyid* clams, *bathymodiolin* mussels, and serpulids—were noted. Additionally, the presence or absence of hard substrate in the form of carbonate rock was noted for each photo. In ArcGIS 10.6.1, these species and substrate presences were resampled into 5 m grid cells in the survey area to include only spatially explicit presence points, and they were mapped over multibeam bathymetric data collected by *AUV Sentry*. The presence of serpulids was categorized as “seep-associated” or “non seep-associated,” where seepage was indicated by the presence of seep-related foundation species found in the same grid cell.

### Statistical Analysis

Quantification and statistical analyses are described in the Results sections and figure legends. Comparisons were performed using ANOVA and statistical significance was declared at P < 0.05. Statistical analyses of beta diversity were performed with Primer E.

## Supporting information

Full Supplement

## Supplementary materials

Fig. S1. Relative abundance of bacterial phylotypes, based on 16S rRNA.

Fig. S2. Phylogenetic relationships of the dominant 16S rRNA and pmoA sequences recovered from two polychaete species featured in this study.

Fig. S3. Fluorescence Microscopy of *Bispira* sp. and *Laminatubus* sp. crown radioles.

Fig. S4. FISH and TEM Microscopy of *Bispira* sp. and *Laminatubus* sp. crown radioles.

Fig. S5. Fluorescence Microscopy of *Bispira* sp. and *Laminatubus* sp. whole specimens.

Fig. S6. ^13^C-labelled dissolved inorganic carbon generated during shipboard experiments.

Table S1: Sample locations along the west coast of Costa Rica, along with dive information and sampling date

Table S2: δ^13^C (‰) of various body tissues for both species, including native worms and 16-month transplants

Table S3: Initial conditions of labeled ^13^CH_4_ at the start of each isotope incubation experiment.

Data File S1: MiSeq 16S rRNA sequence data collected in this study

## Acknowledgements

We thank the captains and crew of the *RV Atlantis, HOV Alvin* pilots and technicians, as well as scientific participants of AT 37-13 and AT 42-03, especially O. Pereira, L. McCormick, C. Seid, and A. Durkin for their assistance at sea. We also thank the captain and crew of the *RV Falkor, ROV SuBastian* pilots and technicians, as well as scientific participants of FK190106, as well as J. Gonzalez, A. Crémière, J. Magyar, and S. Connon for their assistance and contributions to the isotope, dissolved IC, nanoSIMS, and microbial community analysis, respectively. C. Roman (University of Rhode Island, as co-PI) and J. Cortes Nunez (University of Costa Rica, international collaborator) played significant roles as co-PI’s during both expeditions. Additionally, Occidental College undergraduates M. Cazin, K. Ruis, C. Brzechffa assisted with microbial community and microscopy analysis. Support was provided by a Faculty Enrichment Grant through Occidental College. The research was mainly supported by the US National Science Foundation grants OCE 1635219 (EC), OCE 1634172 (LL & GR), 1634002 (VO).

## Author contributions

S.G. conducted DNA analysis, including 16S rRNA barcoding, fluorescent microscope analyses, analyzed experimental data, wrote the manuscript with input from co-authors, and participated in both expeditions. E.T. performed electron microscopy analyses and participated in AT42-03. A.K. performed Sentry data analysis and wrote the manuscript. K.D. designed and performed the incubation experiments and participated in both expeditions. S.M. designed and performed the incubation experiments and participated in both expeditions. F.W. performed fluorescent microscope analyses. R.L performed the isotope analyses. G.R. was a principal investigator on the NSF-funded project, fixed the specimens for study (including seamount worms), coordinated and interpreted the electron microscope analyses, identified and is naming the worm species, and participated in all expeditions. L.L. was a principal investigator on the NSF-funded project, coordinated isotope analyses, wrote the manuscript, and participated in both expeditions. E.C. was a principal investigator on the NSF-funded project, and Chief Scientist on both expeditions. V.O. was a principal investigator on the NSF-funded project, designed the incubation experiments, and participated in both expeditions. All authors contributed to data interpretation and editing of the paper.

## Data Availability

The raw barcode sequence data and QIIME processed data are available in DataFile S1. 16S rRNA sequences for bacterial isolates are available from GenBank under accession numbers TBD – submitted July 24, 2019. Additionally, the sequences reported in this paper will be deposited in National Center for Biotechnology Information database through Sequence Read Archive under BioProject (accession no. <u>TBD</u>).Animal images and specimens were vouchered (*Laminatubus* cat #A9589 and *Bispira* cat #A9598) for long-term archiving into the Benthic Invertebrate Collection at Scripps Institution of Oceanography (https://sioapps.ucsd.edu/collections/bi/).

