## Supplementary material for "Sequestration of Methane by Symbiotic Deep-Sea Annelids: Ancient and Future Implications of Redefining the Seep Influence": Full Supplement

**Supplemental Information – Goffredi et al.**

Fig. S1. Relative abundance of bacterial phylotypes, based on 16S rRNA.
Fig. S2. Phylogenetic relationships of the dominant 16S rRNA and pmoA sequences recovered from two polychaete species featured in this study.
Fig. S3. Fluorescence Microscopy of *Bispira* sp. and *Laminatubus* sp. crown radioles.
Fig. S4. FISH and TEM Microscopy of *Bispira* sp. and *Laminatubus* sp. crown radioles.
Fig. S5. Fluorescence Microscopy of *Bispira* sp. and *Laminatubus* sp. whole specimens.
Fig. S6. $^{13}$C-labelled dissolved inorganic carbon generated during shipboard experiments.
Table S1: Sample locations along the west coast of Costa Rica, along with dive information and sampling date
Table S2: $\delta^{13}$C (‰) of various body tissues for both species, including native worms and 16-month transplants
Table S3: Initial conditions of labeled $^{13}CH_4$ at the start of each isotope incubation experiment.
Data File S1: MiSeq 16S rRNA sequence data collected in this study

**Supplemental Figure Legends**

**Fig. S1 │Relative abundance of bacterial phylotypes, based on 16S rRNA.** Bacterial community structure (to the phylotype level; 99% 16S rRNA sequence similarity) for crown radioles of the serpulid *Laminatubus* n. sp. and sabellid *Bispira* n. sp. from Jaco Scar and Mound 12 seeps, Costa Rica. "Non-seep" specimens were collected from inactive areas at 320-520 m depth. Each color on the graph represents a distinct genus-level phylotype or lowest level available. Phylotypes were grouped to 97% 16S rRNA sequence similarity. Dominant phylotypes are indicated in the key. Genera that were not putative aerobic methanotrophs are shown in gray or black. See SI Appendix Table 4-6 for raw and processed data, as well as representative sequences for all dominant hits.

**Fig. S2│Phylogenetic relationships of the dominant 16S rRNA and pmoA sequences recovered from two polychaete species featured in this study.** Targeted PCR amplification of (A) Methylococcales-specific 16S rRNA sequences (663 bp final length sequence) recovered using the 27F/850R primer pair and (B) the particulate methane monooxygenase gene sequence (pmoA; 408 bp final length sequence), recovered using the 189f/661r primer pair, from the radioles of the serpulid *Laminatubus* n. sp. and *Bispira* n. sp, as well as surrounding seawater samples taken by CTD above the seep sites (all shown in bold). Close relatives were assigned using BLAST. Each tree is based on neighbor-joining analysis, constructed with a Jukes-Cantor distance model, and the resulting tree topology was evaluated by bootstrap analysis (* at nodes indicate bootstrap support >60, from 1000 resampled data sets). The scale bars represent % substitutions per site. HTV = hydrothermal vent.

**Fig. S3 | Fluorescence Microscopy of *Bispira* sp. and *Laminatubus* sp. crown radioles.** (A-B) Fluorescence microscopy of annelid radioles showing comparison between the general bacterial probe Eub338 I-III labelled with Cy3 (A, D) and probe MTC851 labelled with FITC, designed in this study to be an exact match to the MOX symbionts of both *Laminatubus* and *Bispira* from Jaco Scar, plus the overlay (C,F). *Bispira* n. sp. is shown in A-C. *Laminatubus* n. sp. is shown in D-F. DAPI-stained nuclei of host cells are shown in blue. All scale bars are 20 μm.

**Fig. S4 | Microscopy of *Bispira* sp. and *Laminatubus* sp. crown radioles.** Fluorescence microscopy of (A) a mannose-specific biofilm, via HHA staining (in red) surrounding the radiolar crown tip of *Bispira*. DNA of host cell nuclei and nearby bacterial symbionts shown via DAPI-staining in cyan. Fluorescence microscopy of (B-C) lipid-rich organelles, via FM46-4 staining (in red) surrounding the radiolar tips of both *Bispira* and *Laminatubus*, respectively. A Methylococcales/Marine Methylotrophic Group 2 specific FISH probe (MTC851) is shown in green, and DNA is shown in blue/cyan via DAPI counterstain. Transmission electron microscopy of *Laminatubus* n. sp. (D-E) and *Bispira* n. sp. (F-G) radioles, showing MOX bacteria, with dense internal membranes (asterisks), completely engulfed by host cells (arrowheads). A-C scale bars are 10 μm, D-G scale bars are 500 nm.

**Fig. S5 | Fluorescence Microscopy of *Bispira* sp. and *Laminatubus* sp. whole specimens.** For both species, the digestive tract is a straight, ciliated tube, with a simple foregut, a stomach, intestine, and hindgut. The anterior portion of the gut is constricted by mesenteries, and expands into a series of spherical chambers. *Bispira* whole specimen (A), 3-μm section of specimen embedded in Steedman's resin (B). Letters C-E correspond to FISH images of various digestive tract regions, including the mouth (C), intestine (D), and hindgut (E). A Methylococcales/ Marine Methylotrophic Group 2 specific FISH probe (MTC851) is shown in green (none are positive), and host nuclei are shown in blue via DAPI counterstain. *Laminatubus* whole specimen (F), 3-μm section of specimen embedded in Steedman's resin (G). Letters H-J correspond to FISH images of various body regions, including the radioles (H), mouth (I), and intestine (J). A general bacterial probe set Eub338 I-III labelled with Cy3 is shown in orange (only the radioles are positive), and host nuclei are shown in blue via DAPI counterstain. Inset in J shows non-bacterial autofluorescent objects within a region of the intestine. All scale bars are 100 μm.

**Fig. S6 | $^{13}$C-labelled dissolved inorganic carbon generated during shipboard experiments.** The generation of substantial $^{13}$C-labelled dissolved inorganic carbon in the surrounding seawater was measured within 15 hours of incubation for both annelid species (*Bispira* and *Laminatubus*), confirming $CH_4$ conversion to $CO_2$ by the MOX bacterial symbionts. $^{13}$C-labelled methane ranged from 25 at% (*), to 50-55at%, to 100 at% (**). See SI Appendix Table 3 for more details.

**Supp Table 1**: Sample locations along the west coast of Costa Rica, along with dive information and sampling date.

| Site | Geo location | Date | Dive #[1] | Depth (m) |
|---|---|---|---|---|
| Jaco Scar – Active Seep | 9.11715°N / 84.84131°W | Oct 17-18, 2018<br>Nov 4, 2018 | AD4971-72<br>AD4989 | 1824 |
| JS 600m away – inactive | 9.11730°N / 84.83961°W | Oct 17, 2018 | AD4971 | 1796 |
| JS 1400m away – inactive | 9.11491°N / 84.83972°W | Oct 19, 2018 | AD4973 | 1887 |
| Mound 12 – Active Seep | 8.93075°N / 84.3128°W | May 21, 2017<br>Oct 24, 2018 | AD4906<br>AD4978 | 995<br>999 |
| Seamount 6 - inactive | 7.68025°N / 85.91171°W | Jan 22, 2019 | SO227 | 527 |
| Coco South - inactive | 5.46692°N / 87.13112°W | Jan 19, 2019 | SO224 | 321 |

[1] AD = DSRV *Alvin* dive number (Woods Hole Oceanographic Institute). SO = ROV *SubBastian* dive number (Schmidt Ocean Institute).

**Supp Table 2**: $\delta^{13}C$ (‰) of various body tissues for both species, including native worms and 16-month transplants (avg ± 1 SD).

| | $\delta^{13}C$ (‰) | | |
|---|---|---|---|
| **Tissue region** | ***Bispira* (native)** | ***Laminatubus* (native)** | ***Laminatubus* (transplanted)** |
| crown<br>body<br>gut | -50.5 ± 1.3<br>-47.8 ± 0.9<br>-48.0 ± 1.4<br>(n = 2) | -56.9 ± 0.9<br>-56.9 ± 0.5<br>-57.9 ± 0.6<br>(n = 4) | -50.5 ± 0.7<br>-51.5 ± 0.5<br>-51.9 ± 0.8<br>(n = 6) |

**Supp Table 3**: Initial conditions of labeled $^{13}CH_4$ (concentration and the $^{13}C$ atom percent; atom%) at the start of each isotope incubation experiment.

| Incub # | Time Incubated | Sample type | Vol SW (mL) | Methane Conc. (mM) | $^{13}CH_4$ At% |
|---|---|---|---|---|---|
| 10854-1-5 | 24-37 h | *Bispira* without tubes | 40 | 1.11 | 50 |
| 10854-6-8 | 69 h | *Laminatubus* without tubes | 40 | 1.11 | 50 |
| 10854-A | 80 h | *Laminatubus* on carbonate | 1400 | 0.22 | 55 |
| 10861 | 105 h | *Laminatubus* on carbonate | 450 | 0.37 | 25 |
| 10882 | 35 h | *Bispira* on carbonate | 600 | 0.32 | 25 |
| 11669 | 24 h | *Bispira* on Lamellibrachia tubes | 800 | 0.11 | 100 |

**Data File 1**: Excel file of raw MiSeq 16S rRNA sequence data collected in this study (sheet 1), normalized and trimmed MiSeq 16S rRNA sequence data collected in this study (sheet 2), and representative Methylococcales - Marine Methylotrophic Group 2 sequences associated with *Laminatubus*, *Bispira*, and water samples from Jaco Scar, Costa Rica (sheet 3). Attached as separate .csv file

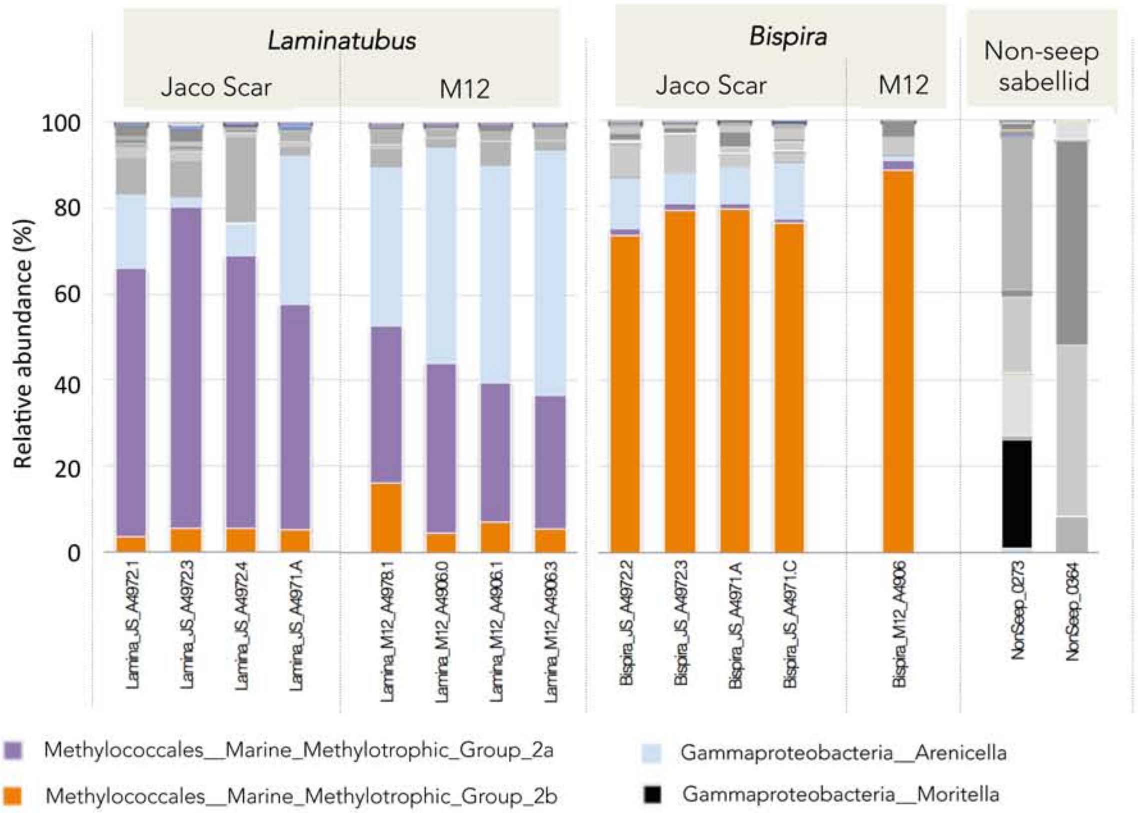


Fig. S1: Relative abundance of bacterial phylotypes, based on 16S rRNA. Bacterial community structure (to the phylotype level; 99% 16S rRNA sequence similarity) for crown radioles of the serpulid Laminatubus n. sp. and sabellid Bispira n. sp. from Jaco Scar and Mound 12 seeps, Costa Rica. “Non-seep” specimens were collected from inactive areas at 320-520 m depth. Each color on the graph represents a distinct genus-level phylotype or lowest level available. Phylotypes were grouped to 97% 16S rRNA sequence similarity. Dominant phylotypes are indicated in the key. Genera that were not putative aerobic methanotrophs are shown in gray or black. See SI Appendix Table 4-6 for raw and processed data, as well as representative sequences for all dominant hits.

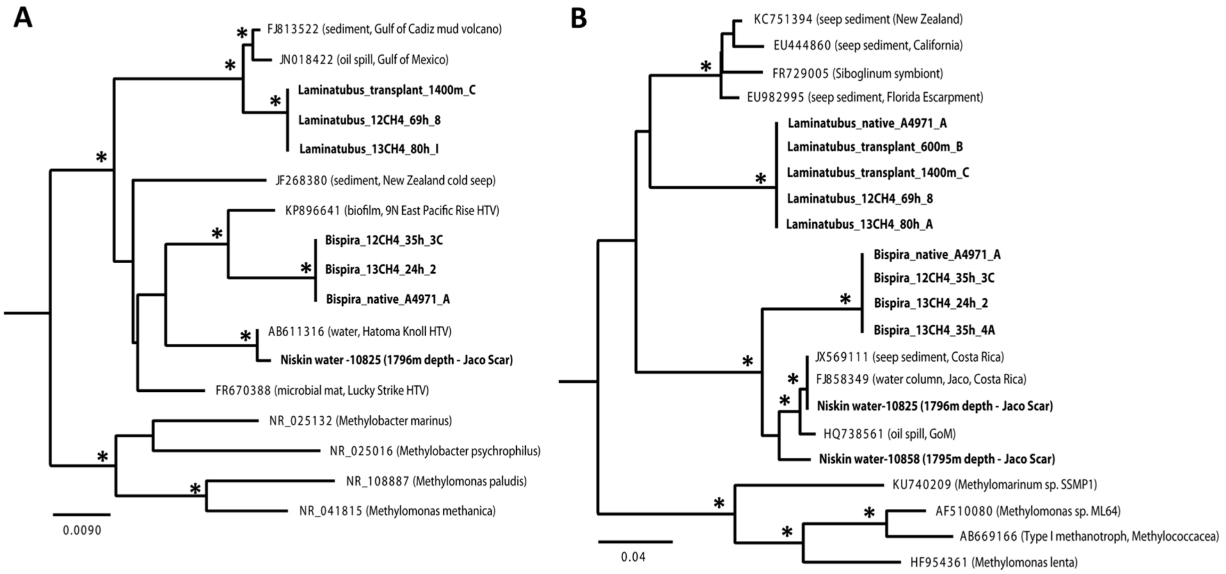


Fig. S2. Phylogenetic relationships of the dominant 16S rRNA and pmoA sequences recovered from two polychaete species featured in this study. Targeted PCR amplification of (A) Methylococcales-specific 16S rRNA sequences (663 bp final length sequence) recovered using the 27F/850R primer pair and (B) the particulate methane monooxygenase gene sequence (pmoA; 408 bp final length sequence), recovered using the 189f/661r primer pair, from the radioles of the serpulid Laminatubus n. sp. and Bispira n. sp, as well as surrounding seawater samples taken by CTD above the seep sites (all shown in bold). Close relatives were assigned using BLAST. Each tree is based on neighbor-joining analysis, constructed with a Jukes-Cantor distance model, and the resulting tree topology was evaluated by bootstrap analysis (* at nodes indicate bootstrap support >60, from 1000 resampled data sets). The scale bars represent % substitutions per site. HTV = hydrothermal vent.

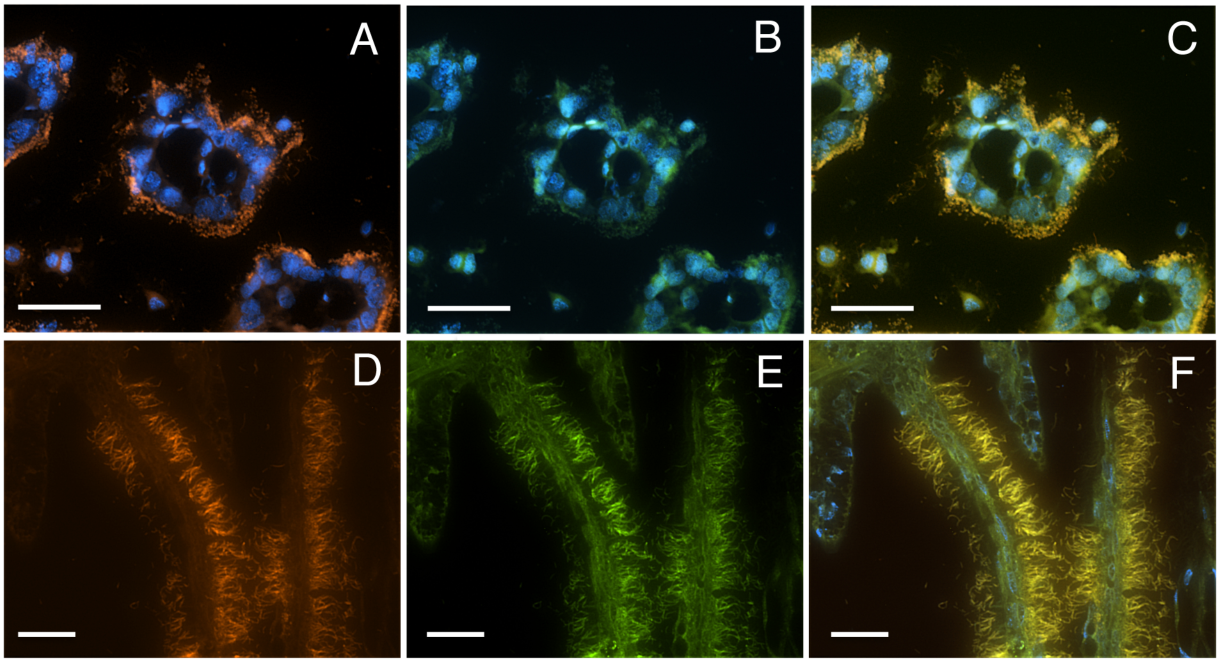


Fig. S3 Fluorescence Microscopy of Bispira sp. and Laminatubus sp. crown radioles. (A-B) Fluorescence microscopy of annelid radioles showing comparison between the general bacterial probe Eub338 I-III labelled with Cy3 (A, D) and probe MTC851 labelled with FITC, designed in this study to be an exact match to the MOX symbionts of both Laminatubus and Bispira from Jaco Scar, plus the overlay (C,F). Bispira n. sp. is shown in A-C. Laminatubus n. sp. is shown in D-F. DAPI-stained nuclei of host cells are shown in blue. All scale bars are 20 μm.

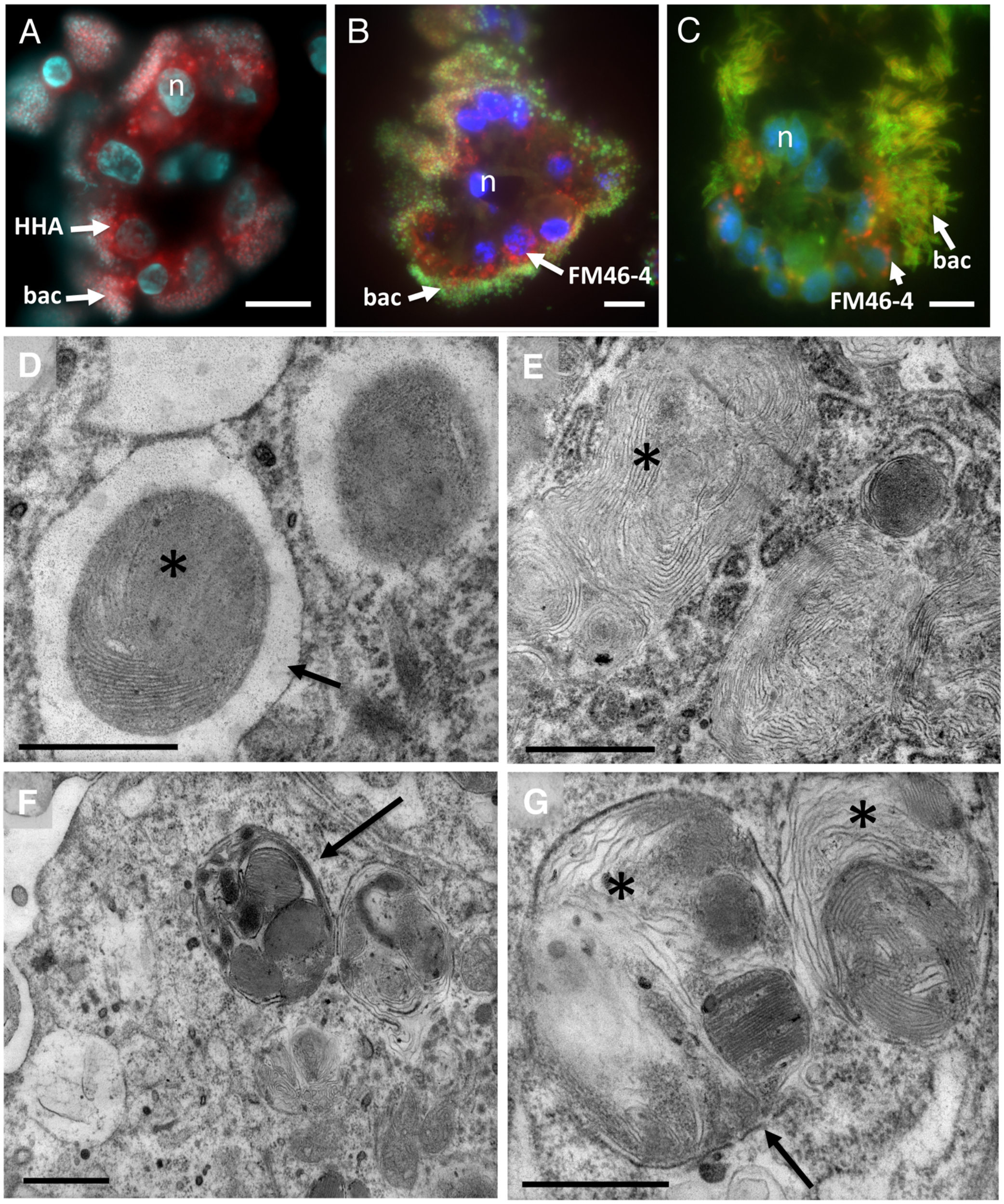


Fig. S4. Microscopy of Bispira sp. and Laminatubus sp. crown radioles. Fluorescence microscopy of (A) a mannose-specific biofilm, via HHA staining (in red) surrounding the radiolar crown tip of Bispira. DNA of worm host cell nuclei and nearby bacterial symbionts shown via DAPI-staining in cyan. Fluorescence microscopy of (B-C) lipid-rich organelles, via FM46-4 staining (in red) surrounding the radiolar tips of both Bispira and Laminatubus, respectively. A Methylococcales/Marine Methylotrophic Group 2 specific FISH probe (MTC851) is shown in green, and DNA is shown in blue/cyan via DAPI counterstain. Transmission electron microscopy of Laminatubus n. sp. (D-E) and Bispira n. sp. (F-G) radioles, showing MOX bacteria, with dense internal membranes (asterisks), completely engulfed by host cells (arrowheads). A-C scale bars are 10 µm, D-G scale bars are 500 nm.

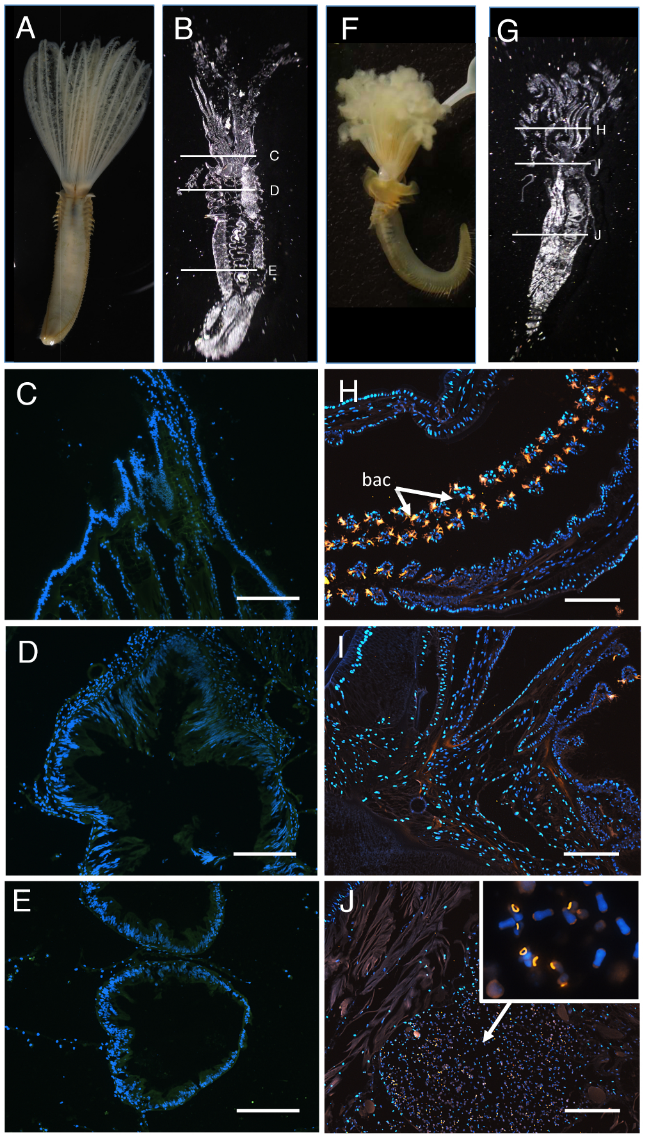


Fig. S5. Fluorescence Microscopy of Bispira sp. and Laminatubus sp. whole specimens. For both species, the digestive tract is a straight, ciliated tube, with a simple foregut, a stomach, intestine, and hindgut. The anterior portion of the gut is constricted by mesenteries, and expands into a series of spherical chambers. Bispira whole specimen (A), 3-μm section of specimen embedded in Steedman's resin (B). Letters C-E correspond to FISH images of various digestive tract regions, including the mouth (C), intestine (D), and hindgut (E). A Methylococcales/ Marine Methylotrophic Group 2 specific FISH probe (MTC851) is shown in green (none are positive), and host nuclei are shown in blue via DAPI counterstain. Laminatubus whole specimen (F), 3-μm section of specimen embedded in Steedman's resin (G). Letters H-J correspond to FISH images of various body regions, including the radioles (H), mouth (I), and intestine (J). A general bacterial probe set Eub338 I-III labelled with Cy3 is shown in orange (only the radioles are positive), and host nuclei are shown in blue via DAPI counterstain. Inset in J shows non-bacterial autofluorescent objects within the intestine. Scale bars are 100 μm.

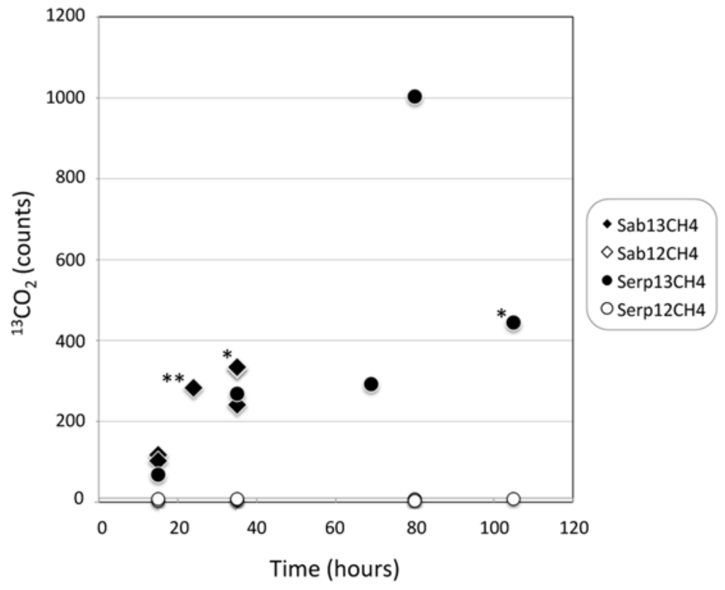


Fig. S6: 13C-labelled dissolved inorganic carbon generated during shipboard experiments. The generation of substantial 13C-labelled dissolved inorganic carbon in the surrounding seawater was measured within 15 hours of incubation for both annelid species (Bispira and Laminatubus), confirming CH4 conversion to CO2 by the MOX bacterial symbionts. 13C-labelled methane ranged from 25 at% (*), to 50-55at%, to 100 at% (**).
See SI Appendix Table 3 for more details.
